# Identification of Phosphotyrosine-Mutant Desmin in Human Pancreatic Cancer

**DOI:** 10.1101/2024.10.14.618246

**Authors:** Nancy Kendrick, Costel C. Darie, Matt Hoelter, Andrew Koll, Jon Johansen

## Abstract

Pancreatic ductal adenocarcinoma (PDAC), a deadly cancer with a 5-year survival rate of ∼12%, is characterized by frequently mutated KRAS, early metastasis, and extensive desmoplasia. The latter, a formation of dense fibrotic tissue generated by pancreatic stellar cells (PSC) and tumor cells, makes up to 80% of the tumor mass and leads to treatment failure. To search for novel protein biomarkers for actionable tyrosine kinases, we used a combination of two orthogonal protein analysis methods, western blotting (WB) and mass spectrometry (MS). That is, we used 1D/2D phosphotyrosine (pTyr) WB in combination with nano liquid chromatography tandem mass spectrometry (NanoLC-MS/MS) to analyze homogenates of 6 PDAC tumor samples and 5 normal adjacent tissue controls. Surprisingly, we found a novel, abundant 55 kDa pTyr-protein in 2/6 tumor samples and identified it as mutated pTyr-desmin. Further proteomics analysis of the purified protein cut from multiple Coomassie-stained 2D gels revealed that the mutant amino acid, tyrosine (D399Y), is phosphorylated (pTyr). A possible role for mutant pTyr-desmin in PDAC metastasis is discussed, along with peptide inhibitor drugs.

## Introduction

Pancreatic ductal adenocarcinoma (PDAC) is a deadly cancer characterized by a high percentage of KRAS mutations, extensive desmoplasia, and rapid metastasis with a 5-year survival rate of 12% [1, 2]. Desmoplasia is a formation of dense fibrotic tissue containing myofibroblasts, pancreatic stellar cells (PSC), and tumor cells comprising 50 - 80% by volume of the PDAC tumor. In most patients it probably begins when mutant KRAS in tumor cells triggers the release of cytokines, chemokines, and growth factors that activate nearby quiescent PSCs causing them to release collagens and other stromal proteins [3]. Yes-associated protein (YAP), a transcriptional co-activator and effector of the Hippo signaling pathway, is also involved [4] as are Src Family Tyrosine Kinases (SFKs) [5]. Interactions between PSC and tumor cells exacerbate desmoplasia to the point where high interstitial pressure causes vascular collapse that prevents drug entry [3]. Thus, desmoplasia itself is a PDAC drug target [6].

Our laboratory has shown that phosphotyrosine western blotting (pTyr WB) is sensitive enough to detect activated receptor tyrosine kinase (pTyr-RTK) driver proteins in lung squamous cell carcinoma [7, 8]. Since SFKs, KRAS, and YAP are involved in PDAC, and since wild type pTyr-RTKs might be unknown drivers, we decided to use WB to probe for these proteins in PDAC tumor tissues purchased from a biobank. If a promising pTyr-RTK driver or substrate for a SFK driver of PDAC were found, then, after verification, an antibody against the pTyr-protein biomarker could be used for immunohistochemistry (IHC) testing of human biopsies to help guide drug treatment. Many TK inhibitors are now available [9].

We began by comparing WB patterns of six PDAC tumor and five normal adjacent tissue (NAT) samples using antibodies against KRAS, YAP, Src, and pTyr-protein. Two of the six PDAC tumors [P1 and P2] [and none of the rest] showed abundant pTyr-protein bands at about 60 kDa as well as dark KRAS and YAP1 bands relative to NAT controls. To identify the ∼60 kDa pTyr-protein(s), we performed 2D pTyr WB of the P1 and P2 samples with matching 2D Coomassie stained gels to obtain enough purified stained protein for analysis by mass spectrometry (MS). The ∼60 kDa protein was identified as pTyr-mutant desmin with the mutation being aspartate 399 mutated to tyrosine 399 (D399Y). The pTyr residue was identified as mutant Y399. Possible consequences and ramifications of this surprising and potentially important observation are discussed.

## Results

All tumor and NAT tissues samples were dissolved in SDS buffer with protease and phosphatase inhibitors as described in methods to ensure complete protein recovery. All MS was performed on 1D bands or 2D spots cut from sodium dodecylsulfate polyacrylamide electrophoresis (SDS PAGE) gels [10].

Figure 1 shows KRAS, YAP, Src, and pTyr WB patterns from four 1D gels identically loaded with 40 µg total protein/lane from five alternating PDAC tumor/NAT samples and a singleton tumor sample, P0-T, from an earlier date. Samples P1 and P2 showed patterns similar to each other and different from the rest.

**Figure 1.**
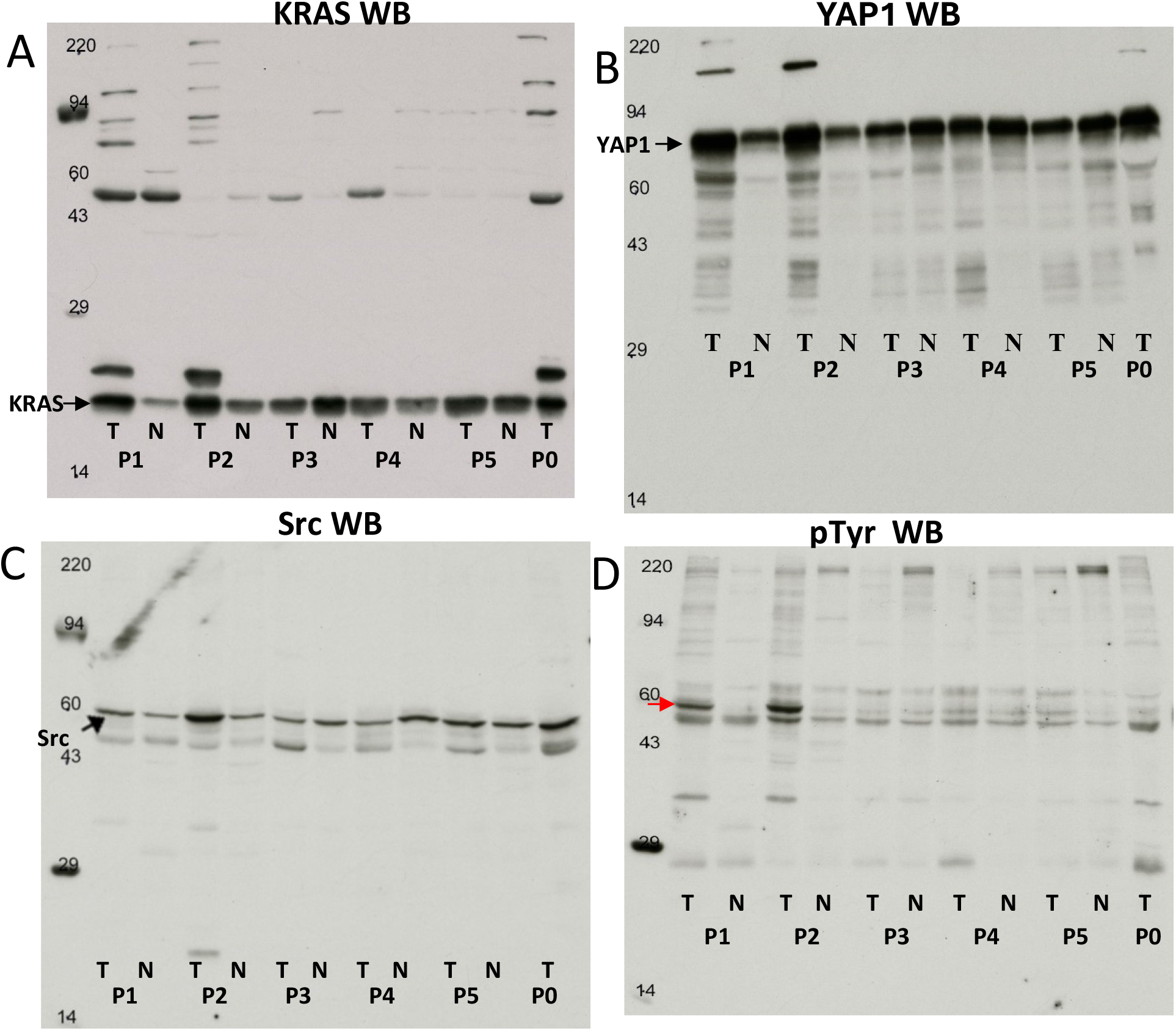
KRAS, YAP, Src and pTyr western blots of six pancreatic tumor (P1-P5 & P0) and five normal adjacent tissue samples. Equal protein loading (40 ug/lane) was used for WB normalization [12]. The heavily loaded molecular weight markers are myosin, 220 kDa; phosphorylase A, 94 kDa; catalase, 60 kDa; actin, 43 kDa; carbonic anhydrase, 29 kDa; and lysozyme, 14 kDa. Red arrow indicates an unknown pTyr-protein of interest.

### Summary of Results for Figure 1

- The KRAS WB (Figure 1A) showed the 21 kDa KRAS protein band was strongly expressed in most of the samples and clearly more abundant in P1 and P2 tumors versus controls, in contrast to P3, P4 and P5 tumor samples. (P0 had no control). Samples P1-T, P2-T and P0-T showed several higher MW bands in the KRAS WB including a dark band at ∼24 kDa of unknown identity. We speculated these might be Ras covalent binding partners or Ras nanoclusters [11], but there was not enough material for identification. The latter would require affinity resin purification before MS analysis.
- The YAP1 WB (Figure 1B) showed that this protein was strongly expressed in most of the samples and clearly more abundant in P1 and P2 tumors versus controls.
- The 60 kDa Src WB bands (Figure 1C) were a little darker in the P1 and P2 tumor samples versus NAT controls but not dramatically so. The bands were roughly the same intensity for P3, P4 and P5 tumor versus controls.
- The pTyr WB (Figure 1D) showed a relatively dark pTyr protein band near the 60 kDa marker for P1 and P2 tumor samples that was absent from both controls (red arrow).

Since tyrosine phosphorylation is important for many cancer cell processes [12, 13], we proceeded with identification of the unknown ∼60 kDa pTyr-protein using purified protein cut from 2D gels.

The next round of analysis employed SDS-compatible 2D electrophoresis (2D) [10] in combination with pTyr WB and Coomassie staining of samples P1 and P2. Figure 2 shows 2D pTyr WBs of P1-Tumor (Fig. 2A), P1-NAT (Fig 2B), P2-Tumor (Fig 2C), and P2-NAT (Fig. 2D).

**Figure 2.**
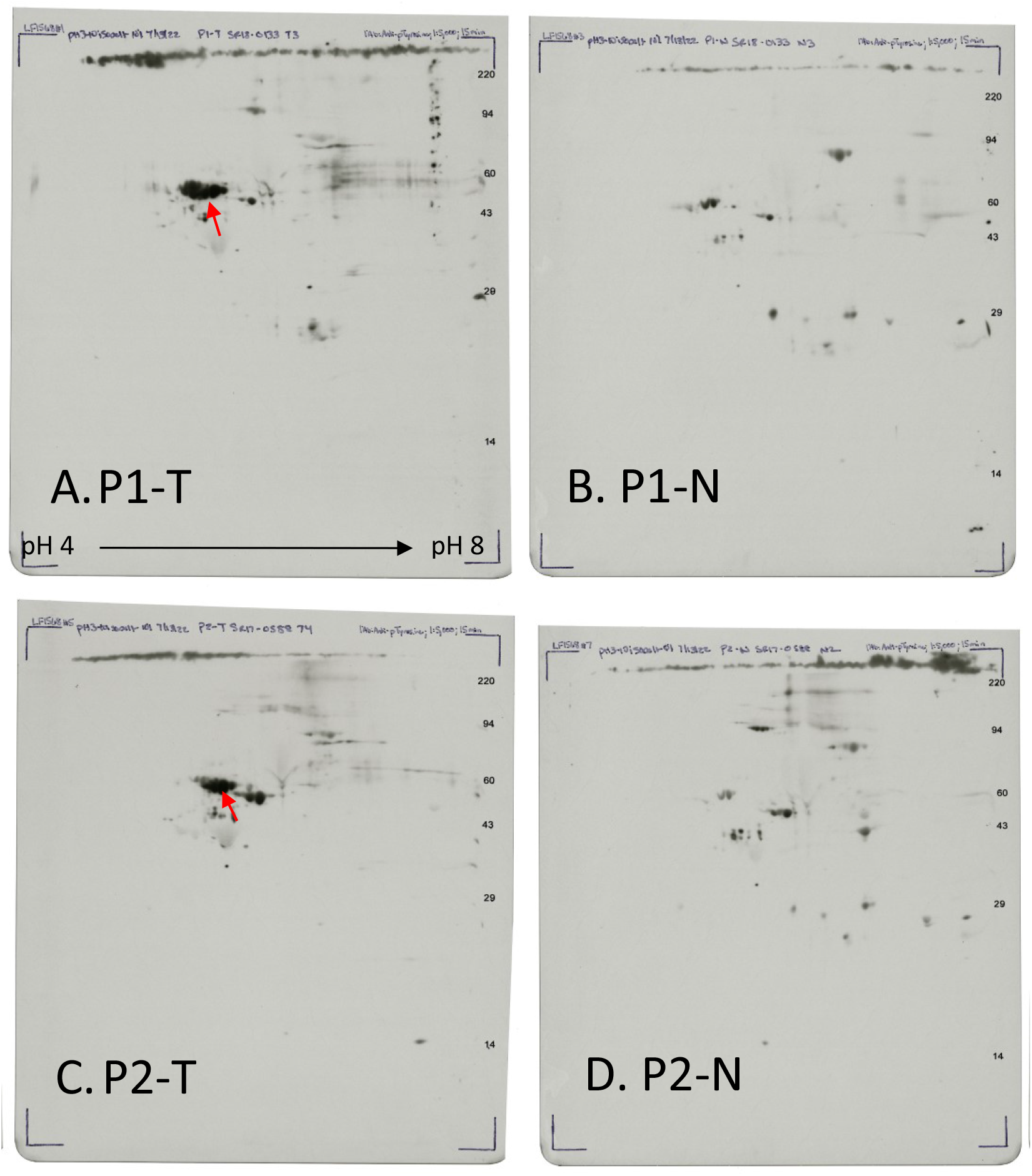
2D pTyr WB patterns of pancreatic tumors P1-T and P2-T along with normal adjacent tissue (NAT) P1-N and P2-N. Red arrows mark the ∼ 60 kDa pTyr-protein in P1-T and P2-T after 15 min film exposure.

Figure 3 shows matching Coomassie stained total protein patterns of these two samples and their controls. Red arrows mark the ∼60 kDa pTyr protein of interest, when present. Note that basic proteins with isoelectric points >8, are not visible on these 2D gels.

**Figure 3.**
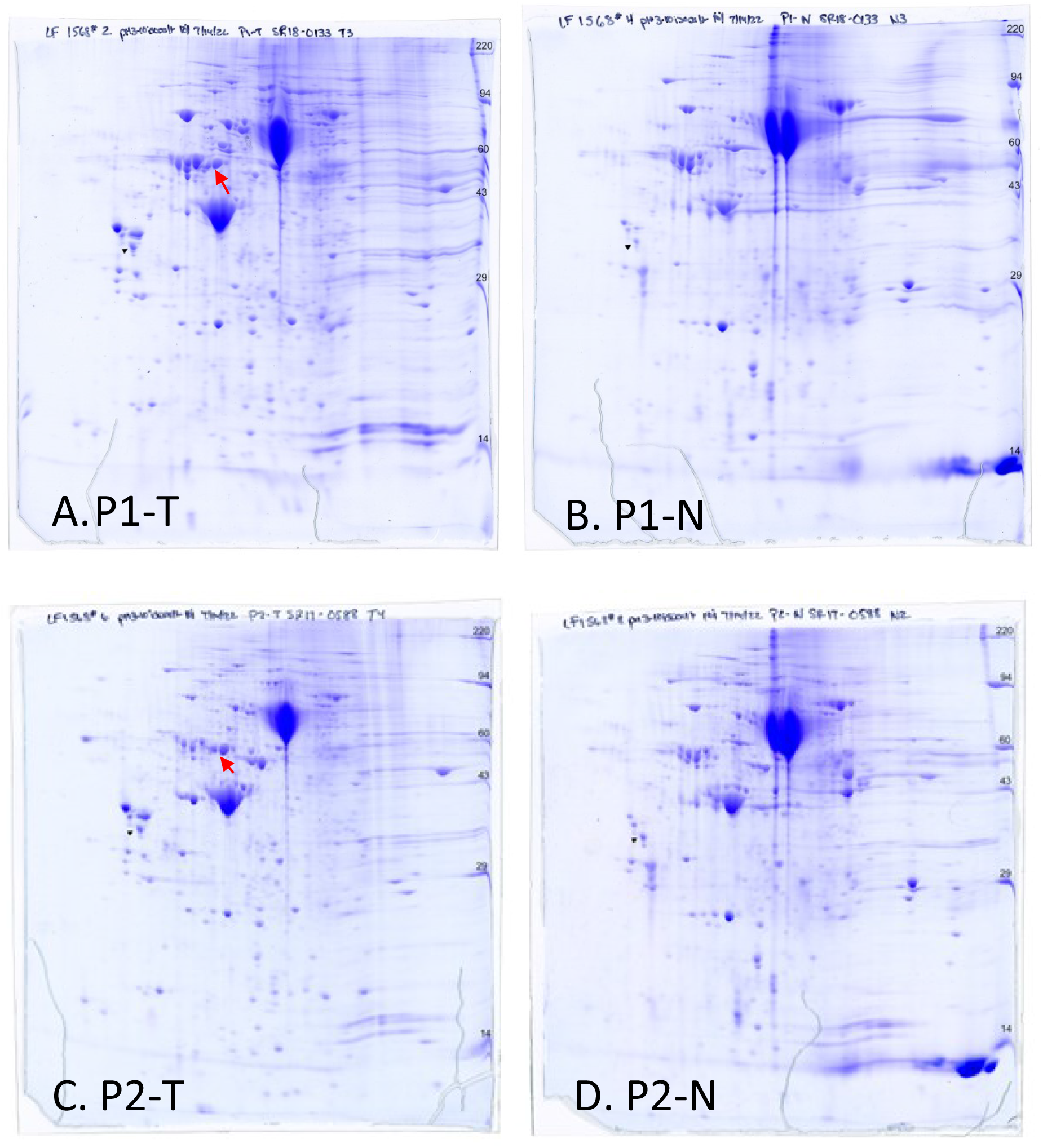
Coomassie stained 2D gel patterns of pancreatic tumors P1-T (A) and P2-T (C), along with NAT tissue patterns P1-N (B) and P2-N (D). LF1568#2, 4, 6, 8. Arrows mark a pTyr protein with four charge-shift isoforms around 60 kDa that were substantially increased in the P1-T and P2-T tumor samples versus their NAT controls.

Figure 4 shows a closeup of the ∼60 kDa pTyr-protein(s) first identified by 1D pTyr WB in Figure 1, then resolved by 2D pTyr WB in Figure 2 and by 2D Coomassie blue staining in Figure 3. The middle overlay images of WB over Coomassie were used to guide Coomassie spot cutting for MS. The ∼60 kDa pTyr unknown protein resolves as a cluster of 4 isoforms (arrows) which are missing or very faint in the NAT samples.

**Figure 4.**
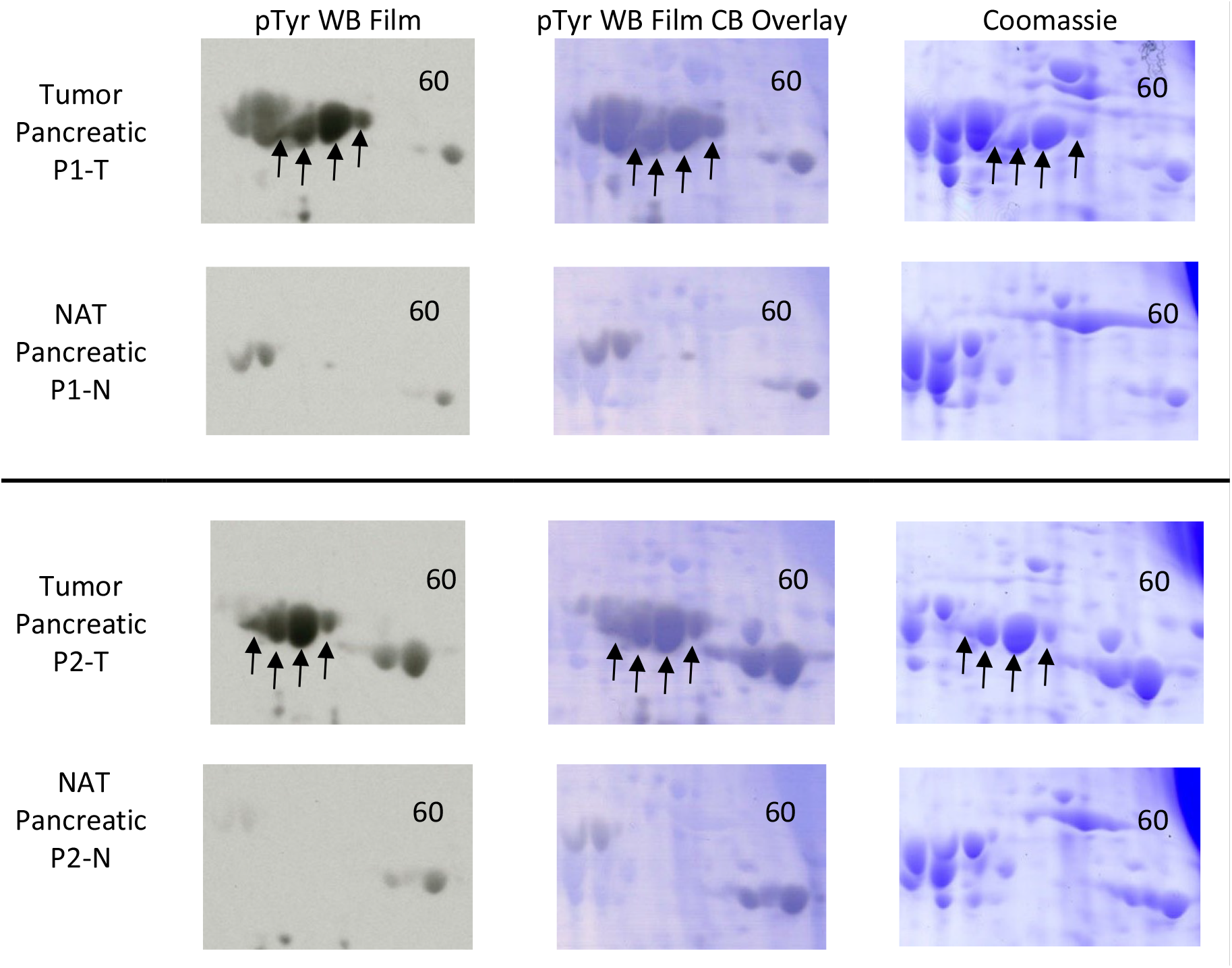
Blowup of 2D gel area of interest showing the pTyr WB film image on the left. The Coomassie blue image on the right, and Coomassie/WB overlay image spot in the center. The latter was used to cut spots from the Coomassie gel for identification by MS.

To determine if the four protein spots were different proteins or isoforms of the same protein, we first cut out each of the four spots from a single Coomassie stained 2D gel for individual identification by NanoLC-MS-MS. All the isoforms were identified as mutant desmin, MW 55 kDa. Charge isoform clusters such as this are common on 2D gels and likely due to post-translational modifications such as cysteine oxidation, carbamylation, glycosylation, or phosphorylation that may change the isoelectric point of a protein [14, 15].

Since the pTyr post-translational modification (PTM) is rare and often difficult to identify [16], we used 2DE to extract purified desmin protein from a relatively large amount of tumor homogenate to identify the phosphorylated tyrosine residue as well as the desmin mutation. To do so we ran 500 ug of the P1-T and P2-T samples each on three large-format 2D gels (1500 ug total). We estimated that relative to our internal standard, each pTyr-protein cluster (including 4 isoforms combined) was ≥2 ug protein. Finally, we excised the pTyr-protein 4-spot clusters in Figure 4 from the six 2D gels and combined them (> 12 ug) for analysis by NanoLC-MS/MS [17].

#### The desmin mutation was identified as aspartate 399 (D) to tyrosine 399 (Y) and the phosphorylated amino acid was identified as mutant tyrosine 399 (Y)

NanoLC-MS/MS analysis on a relatively large scale led to a surprising discovery: in addition to wild type desmin [Homo sapiens] (<u>gi|3411132</u>), we also found a mutant desmin [Homo sapiens], gi|71011081. While the canonical desmin has an amino acid sequence MAL**D**VEIATYRK, with aspartate 399 within the amino acid sequence, the mutant protein had the aspartate mutated to tyrosine: MAL**Y**VEIATYR. The peptide from the wildtype desmin protein was identified by the precursor ions with m/z of 470.59 (3+) and 705.41 (2+) that corresponded to peptide MALDVEIATYRK, and by precursor ions with m/z of 475.93 (3+) and 713.41 (2+), which corresponded to peptide mALDVEIATYRK (oxidized Methionine or Mox).

Conversely, the peptide from the mutant desmin was identified by the precursor ions with m/z of 470.59 (3+) and 705.41 (2+) that corresponded to peptide MAL**pY**VEIATYR (pY: phosphorylated tyrosine) and by precursor ions with m/z of 475.93 (3+) and 713.41 (2+), which corresponded to peptide mAL**pY**EIATYR (Mox). The MS and MS/MS spectra that correspond to peptides mAL**D**VEIATYRK from canonical desmin protein and mAL**pY**EIATYRK from the mutant desmin protein are shown in Figures 5 and 6. Unphosphorylated mutant peptide MALYVEIATYR was not detected.

**Figure 5.**
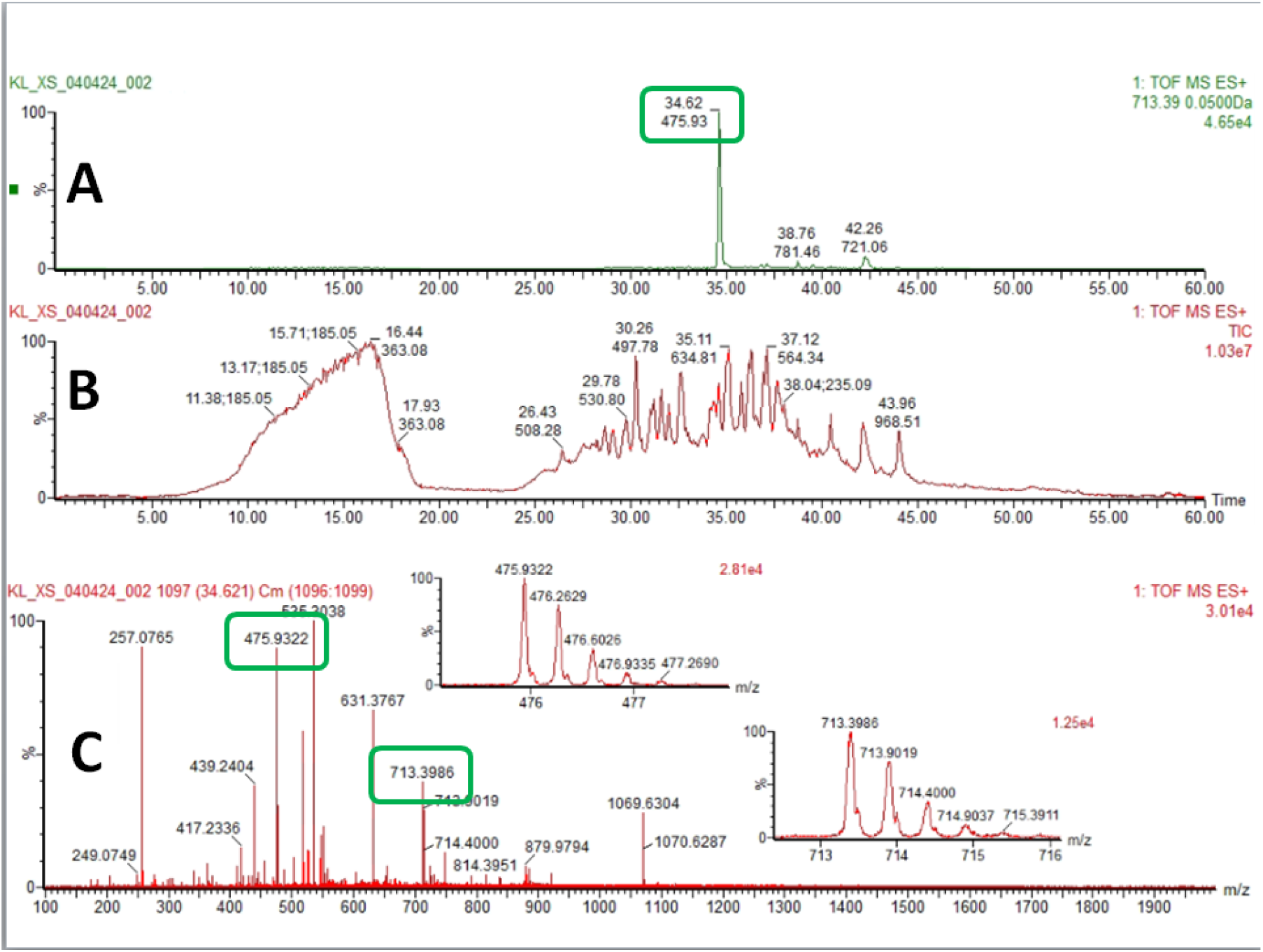
MS and MS/MS spectra that correspond to peptide mAL**D**VEIATYRK A: Extracted Ion Chromatogram (XIC) for the precursor ion with m/z of 475.93(3+) that corresponds to peptides mAL**pY**EIATYR (Mox) and mALDVEIATYRK. B: Total Ion Chromatogram (TIC) from which XIC was obtained. C: MS spectrum in which both (3+) and (2+) precursor ions that correspond to peptides mAL**pY**EIATYR (Mox) and mALDVEIATYRK were identified as precursor ions with m/z of 475.93 (3+) and 713.41 (2+)

**Figure 6.**
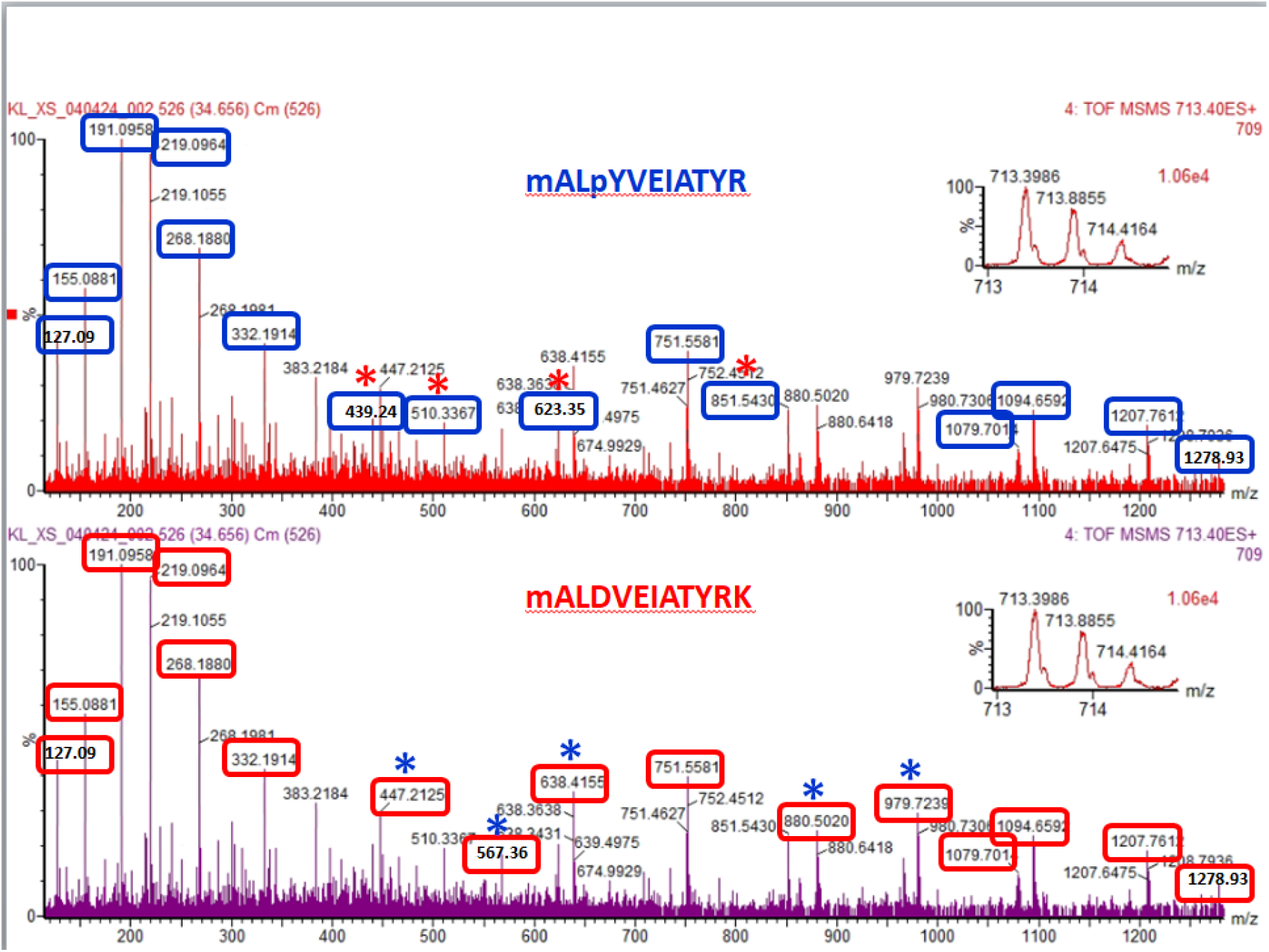
MS/MS spectrum that shows the fragmentation of the precursor ion with m/z 713.40 (2+) that corresponds to peptides mAL**pY**EIATYR (Mox) and mALDVEIATYRK. The fragment ions that correspond to peptide mAL**pY**EIATYR are marked with blue rectangles in the top spectrum. Fragment ions that are unique to this peptide are marked with a red star. The fragment ions that correspond to peptide mAL**D**VEIATYRK are marked with red rectangles in the bottom spectrum. Fragment ions that are unique to this peptide are marked with a blue star.

Further data is provided in Supplemental Figures. Mascot database searches that support the identification of the desmin peptides and the novel desmin isoprotein are provided in Supplemental Figure 1. expanded MSMS spectra from Figure 6 are shown in Supplemental Figure 2. expanded MSMS spectra of the precursor ions with m/z of 475.93 (3+) and 713.41 (2+) that correspond to peptide mALDVEIATYRK (Mox) and mALpYEIATYR (Mox) are shown in Supplemental Figure 3.

## Discussion

While considerable knowledge has been gained from genomic [18], transcriptomic [19] and proteogenomic [20] analysis of PDAC tumors, actionable biomarkers for this deadly cancer remain elusive [21]. For example, KRAS is mutated in 85% of PDAC [1]. But in a clinical trial where 38 PDAC patients with KRAS p.G12C-mutations were treated with sotorasib, a KRAS pG12C inhibitor, only 8 patients had an objective response. Median progression-free survival was 4.0 months, median overall survival 6.9 months, 16 patients had adverse events [22].

Given PDAC’s extensive genomic and transcriptomic heterogeneity [23], its lack of known TK drivers, and the multitude of TK approved drugs [9], we decided to screen for actionable TK *protein* biomarkers in human tumor biopsies using sensitive immunodetection in combination with MS. We were looking for RTKs activated by tyrosine phosphorylation [7], as well as protein substrates of Src which controls many cellular processes [24, 25].

### We found an abundant phosphoprotein, pTyr-mutant desmin D399Y, in 2/6 PDAC tumors

We used 1D/2D pTyr WB [8, 10], in combination with nanoLC-MS/MS [17], to screen for pTyr-protein biomarkers in six PDAC human tumor samples and five NAT controls purchased from a biobank. No high MW pTyr-RTKs were enriched in PDAC tumors versus controls. However, this approach did detect a novel, abundant, 55 kDa pTyr-protein in 2/6 tumor samples and none of the NAT controls. The protein was identified by nanoLC-MS/MS as pTyr-mutant desmin where aspartate 399 in wild type desmin, was replaced by tyrosine 399 in mutant desmin. Furthermore, the mutant tyrosine was phosphorylated on tyrosine 399. Since orthogonal protein analysis methods (WB and MS) were used to compliment and cross-confirm each other, this strengthens our conclusions.

Although serine/threonine phosphorylation of desmin by several kinases has been reported [26], and mutant desmin D399Y detected by genome sequence analysis has been reported for a myopathy-affected family [27], to our knowledge this is the first report of pTyr399-mutant desmin in PDAC.

#### This is a perplexing result

Desmin is a marker for activated pancreatic stellate cells (PSCs) [28], albeit a variable marker [3, 29], and not for PDAC tumor cells. PSCs, non-cancerous mesenchymal cells present in tumor stroma [30], do not have the array of mutated tumor suppressor genes, p53 for example, that lead to the genomic instability of cancer cells [13]. Thus, while PSCs are known to express wild type desmin [28], which was observed, they are unlikely to express *mutated* desmin. Yet pTyr-mutated desmin was an abundant protein in *two* PDAC tumor samples (Figures 1D, 2-4) and is provocative. What’s going on?

#### Our hypothesis

As reviewed by Hingorani [3], PSCs and PDAC tumor cells interact bidirectionally to generate desmoplasia. Mutant KRAS in tumor cells triggers the release of cytokines, chemokines, and growth factors that activate quiescent PSCs [31]. In turn the activated PSCs release extracellular vesicles (EVs) containing miR-21 that stimulate metastasis of the tumor cells [32-34].

We hypothesize that the microRNA-21 stimulates *both* PDAC cancer cells and PSCs to express desmin, and that an unknown amount of the latter in cancer cells is mutant D399A. Since we did not detect any unphosphorylated mutant desmin peptide, mALYEIATYRK, we further hypothesize that a SFK member immediately phosphorylated the mutant tyrosine. Although Src protein expression was only weakly correlated with the pTyr-mutant desmin (Figure 1C), SFKs may still be involved since regulation of their activity is complex [35], and since there are nine SFK members of which three (Src, Yes, and Fyn) are likely to be expressed [25]. Given that Yes Associated Protein (YAP1) expression is markedly increased in tumors 1 and 2 versus their controls (Figure 1B), the Yes kinase is a possible candidate.

### The Clinical Proteomic Tumor Analysis Consortium (CPTAC) did not find pTyr-mutant desmin in their proteogenomics analysis of PDAC tumors [20]. How so?

The words “desmin”, “phosphotyrosine” and “pY” were not found in a text search of the CPTAC manuscript. pTyr-mutant desmin was either not detected by this group or was considered an unimportant passenger mutation. Thus, the abundant pTyr-mutant desmin that we found might be due to a passenger hotspot mutation [36] which has no effect on PDAC growth or metastasis.

Another more likely explanation in our opinion stems from differences in sample preparation. Since our system is compatible with SDS, our procedure (see Methods) includes homogenizing the tumor tissue in SDS buffer containing protease and phosphatase inhibitors along with nucleases. After homogenization, the tube is placed in a boiling water bath for five min to maximize protein solubilization. In many cases the homogenate clarifies. If not, the tubes are centrifuged to remove a small pellet of “cell debris”.

SDS is incompatible with MS instrumentation and must be either omitted or removed [37] from samples prior to MS analysis. Thus, the CPTAC group homogenized PDAC samples in 8M urea without heating followed by centrifugation to remove “cell debris”. It is possible the discarded centrifugation pellet contained undissolved membrane and cytoskeleton proteins which were omitted from their MS analysis. Our method, since it contains SDS and the sample are boiled, most likely solubilized an increased total percentage of the proteins in the tissue samples.

### What advantage might pTyr-mutant desmin provide to tumor cells? Triggering metastasis

Desmin is a type II intermediate filament (IF). IFs are tissue-specific cytoskeleton components that reorganize in response to differentiation related signals [38]. Several IFs have been implicated in cancer epithelial mesenchymal transformation (EMT) and metastasis [39] but details of this complex process remain to be worked out [40]. Kraxner and Koster observed in a reconstituted system that serine/threonine phosphorylation of intermediate filaments (IFs) desmin and vimentin causes disassembly and/or softening of the cell cytoskeleton likely to promote an EMT and cell migration [41].

Allam et al. in reviewing pancreatic stellate cells in PDAC noted: “There is evidence that cancer cells promote their own non-malignant stroma to auto-facilitate their growth and spread: transformed malignant cells can acquire a mesenchymal-like phenotype expressing desmin, vimentin, and α-SMA and can camouflage like resident normal stromal cells [42].” [43].

Kraxner and Koster’s observation provide a mechanism by which IF phosphorylation might promote metastasis. Allam et al.’s statement supports our hypothesis of pTyr-mutant desmin triggering metastasis. If the pTyr-mutant desmin present in PDAC cancer cells imparts a mesenchymal-like phenotype, these cells would be identified as STCs by IHC since desmin is a marker protein for these mesenchymal cells. However, since they are tumor cells expressing a phosphorylated IF protein (desmin) that promotes EMT/metastasis then pTyr-mutant desmin abundance would increase as the metastatic cells multiply. This would explain the relatively high abundance of pTyr-desmin in our samples.

#### *Implications/applications:* A competitive peptide inhibitor drug *might* reduce metastasis

Given PDAC’s extreme heterogeneity, the chain of events outlined here would likely only occur in a tumor subset. We observed pTyr-desmin in 2/6 samples in this report. However, if the hypothesis holds true that 1) SFKs phosphorylate all Y399 mutant desmin and 2) pTyr-mutant desmin triggers metastasis, then the pTyr-desmin protein would be a biomarker for an easily synthesized peptide drug (MAL**Y**VEIATYR) that should interfere with Src phosphorylation of mutant desmin protein. Such a drug would likely have few side effects since it would interfere only with mutant desmin tyrosine phosphorylation and not with wild type desmin’s normal role in muscle.

If the abundance of pTyr-mutant desmin observed here is due to metastatic overexpression of a subclone, then other similar desmin mutations would also be expanded and might serve as biomarkers detectable by IHC.

#### Limitations of the Study

While our identification of pTyr-mutant desmin in 2/6 tumor samples is unequivocal, the small size of this study is a limitation. The pTyr PTM could be present in a small percent of patients or could be an unimportant passenger mutation that has nothing to do with metastasis. While there is evidence in the literature that serine and threonine phosphorylation of intermediate filaments enhances metastasis [41], there is little data on tyrosine phosphorylation. As succinctly summarized by Bracken and Goodall, the EMT is a highly complex process key to tissue and organ development throughout the body. It is controlled by hundreds of transcription factors and microRNAs as well as by phosphorylation events [40]. Unraveling the factors that trigger EMT in different cancers will not be easy.

#### Future Directions

This preliminary study needs to be followed up and verified with a larger number of PDAC tumor samples and NAT controls using pTyr WB and MS. If possible, using research autopsy samples [44], where primary and metastatic tumor samples could be obtained from the same individual, would be useful. If pTyr-mutant desmin is detected in a subset of PDAC tumors and verified to trigger metastasis, then a specific antibody against the mutant desmin pTyr-peptide could be generated for IHC identification of this biomarker in patients.

#### Overall Conclusions and Major Impact

PDAC tumors are very heterogeneous, and their progression is based on stochastic tumor mutations that vary between individuals [23]. Stromal interactions are key as well [3]. Most cancer patients die because of metastatic events, and not due to primary tumors [45]. If we have stumbled onto a novel mechanism of PDAC metastasis, Src phosphorylation of mutated desmin, then a novel treatment becomes theoretically available. Competitive peptide inhibitors targeting the sequence mALYEIATYRK would specifically interfere with SFK phosphorylation of mutant desmin and could be tailored to other desmin mutations if appropriate. Such peptide inhibitors could be synthesized and might have few side effects. Thus, if successful, this might provide a novel path to PDAC treatment.

## Materials and Methods

### Sample Preparation

Five PDAC samples with matched normal adjacent tissue (NAT) were purchased from Spectrum Health Office of Research and Education, Grand Rapids, MI via Accio Biobank Online. A sixth PDAC sample P0, identically prepared, had been purchased in 2011 for another purpose without a NAT control from ILS Bio, LLC, now BioIVT) Chesterfield, MD and stored at −80° C. All tumor samples and controls were received on dry ice and stored at −80° C until sample preparation described below.

Under the definition of human research subjects [45 CFR4 46.102(f)], the Office of Human Resource Protection does not consider research involving only coded private information or biospecimens (retrospective or remnant) to involve human subjects. These specimens were not specifically collected for the proposed research project. Because investigators cannot readily ascertain the identity of the individuals to whom the coded specimens pertain, the requirement for documented informed consent is not applicable.

Sample preparation was performed on ice with ice-cold reagents as follows: Tissue samples were rinsed with ice-cold Tris buffered saline containing 20 mM Tris, 500 mM NaCl, pH 7.5, then placed in a motorized glass-teflon homogenizer. The tissue was homogenized on ice with: Osmotic Lysis Buffer (10 mM Tris, 0.3% SDS) containing protease inhibitors (AEBSF, leupeptin, E-64, 5 mM EDTA, and benzamidine), Phosphatase Inhibitors I and II, RNase, DNase, and 5 mM MgCl_2)_ and then diluted 1:1 with SDS buffer (5% SDS, 10% glycerol, and 62.5 mM Tris, pH 6.8).

After homogenization and dilution, the sample tube was placed in a boiling water bath for 5 min, and the protein concentration was determined by the BCA assay (Pierce/Thermo Fisher). Finally, each sample was diluted to 4 mg/ml with SDS buffer containing 5% beta-mercaptoethanol. Aliquots were stored at −80° C.

## 1D SDS PAGE

SDS slab gel electrophoresis and western blotting was carried out in 10% acrylamide slab gels (13 × 15 cm, 0.75 mm thick), as previously described for the second dimension of 2DE [8]. Electrophoresis was carried out for about 4 hrs at 15 mA/gel. The following proteins (MilliporeSigma) were used as MW standards in one lane on every gel: myosin (220,000), phosphorylase A (94,000), catalase (60,000), actin (43,000), carbonic anhydrase (29,000), and lysozyme (14,000). WB results were normalized by loading constant total protein in lanes as recommended by NIH guidelines [46].

## 2D SDS PAGE

Two dimensional SDS PAGE was performed as previously described [8]. Briefly, isoelectric focusing was performed in polyacrylamide tube gels polymerized with 2% ampholines (130 mm long x 2.3 mm internal diameter) sealed at the bottom with parafilm, were poured using 2% ampholines (Serva Electrophoresis, Heidelberg, Germany). Ampholines were pH 3-10 Iso-Dalt or a 1:1 mixture of pH 4-6 Ampholines and Servalyte pH 5-8. Samples were loaded at the top (basic end) of the polymerized acrylamide tube gel and isoelectric focusing carried out for 9600 volt-hrs.

Tube gels were extruded by air pressure and equilibrated for 10 min in buffer “O” (10% glycerol, 50 mM dithiothreitol, 2.3% SDS and 0.0625 M tris, pH 6.8). The equilibrated tube gels were frozen on dry ice to prevent protein diffusion and thawed immediately before loading onto the 2^nd^ dimension slab gel. Each tube gel was sealed in 1 ml of agarose to the top of a stacking gel overlaying a 10% acrylamide slab gel and electrophoresis carried out for about 4 hr at 15 mA/gel. MW standards were loaded in an agarose well on the basic end of the tube gel. One µg of an IEF internal standard, tropomyosin, was loaded with every sample.

### Western Blotting

After slab gel electrophoresis, gels were placed in transfer buffer (10 mM CAPS, pH 11.0, 10% methanol) and transblotted onto PVDF membranes overnight at 200 mA and approximately 100 volts/ two gels. The blots were stained with Coomassie Brilliant Blue R-250 (Sigma-Aldrich) and scanned.

Coomassie blue stained PVDF membranes were wet in 100% methanol to remove the stain, rinsed in tween-20 (Biorad) TBS (TTBS), and blocked for two hours in 5% non-fat dry milk diluted in TTBS. Primary antibody incubations were performed overnight on an orbital shaker in 2% nonfat dry milk using antibody dilutions shown in the figures. Blots were rinsed 3 × 10 minutes in TTBS and incubated with secondary antibody diluted 1:2000 for 2 hours. Finally, the blots were treated with Pierce ECL reagent (ThermoFisher) and exposed to x-ray film [Kodak BioMax XAR film (ThermoFisher) or GE Amersham Hyperfilm ECL], followed by film development with an automatic Konica Minolta Medical Film Processor SRX-101A.

### Mass spectrometry

Gel spots were digested with trypsin and the resulting peptides mixtures were analyzed by a NanoAcquity UPLC coupled with a QTOF Xevo G2-XS mass spectrometer (both from Waters Corp) according to published procedures [8]. The raw data were converted into mzML files using MS convert and the database search was done against NCBI Human database using propionamide as fixed modification and methionine oxidation and phosphoY/S/T as variable modifications.

## Data Access Statement

The authors confirm that the data supporting the findings of this study are available within the article and its supplementary materials.

