## Supplemental Figures for "Identification of Phosphotyrosine-Mutant Desmin in Human Pancreatic Cancer"

Supplemental FIG. 1A

### MATRIX SCIENCE Mascot Search Results

#### Peptide View

MS/MS Fragmentation of **MALDVEIATYRK**
Found in **gi|181540** in **NCBInr**, desmin [Homo sapiens]

Match to Query 3385: 1424.783448 from(713.399000,2+) scans(3898) rtinseconds(2079.33)
Title: function=4 process=0 scan=526
Data file G:\KL_XS_040424\KL_XS_040424_002.mzML

```
Monoisotopic mass of neutral peptide Mr(calc): 1424.7333
Fixed modifications: Propionamide (C) (apply to specified residues or termini only)
Variable modifications:
M1     : Oxidation (M), with neutral losses 0.0000(shown in table), 63.9983
Ions Score: 54  Expect: 0.17
Matches : 23/140 fragment ions using 39 most intense peaks   (help)
```

| # | b | b++ | b* | b*++ | b0 | b0++ | Seq. | y | y++ | y* | y*++ | y0 | y0++ | # |
|---|---|---|---|---|---|---|---|---|---|---|---|---|---|---|
| 1 | 148.0427 | 74.5250 | | | | | M | | | | | | | 12 |
| 2 | **219.0798** | 110.0435 | | | | | A | ***1278.7052*** | 639.8563 | 1261.6787 | 631.3430 | 1260.6947 | 630.8510 | 11 |
| 3 | **332.1639** | 166.5856 | | | | | L | ***1207.6681*** | 604.3377 | 1190.6416 | 595.8244 | 1189.6576 | 595.3324 | 10 |
| 4 | **447.1908** | 224.0990 | | | 429.1802 | 215.0938 | D | ***1094.5841*** | 547.7957 | 1077.5575 | 539.2824 | 1076.5735 | 538.7904 | 9 |
| 5 | 546.2592 | 273.6332 | | | 528.2486 | 264.6280 | V | ***979.5571*** | 490.2822 | 962.5306 | 481.7689 | 961.5465 | 481.2769 | 8 |
| 6 | 675.3018 | 338.1545 | | | 657.2912 | 329.1493 | E | ***880.4887*** | 440.7480 | 863.4621 | 432.2347 | 862.4781 | 431.7427 | 7 |
| 7 | 788.3859 | 394.6966 | | | 770.3753 | 385.6913 | I | ***751.4461*** | 376.2267 | 734.4196 | 367.7134 | 733.4355 | 367.2214 | 6 |
| 8 | 859.4230 | 430.2151 | | | 841.4124 | 421.2098 | A | ***638.3620*** | 319.6847 | 621.3355 | 311.1714 | 620.3515 | 310.6794 | 5 |
| 9 | 960.4707 | 480.7390 | | | 942.4601 | 471.7337 | T | ***567.3249*** | 284.1661 | 550.2984 | 275.6528 | 549.3144 | 275.1608 | 4 |
| 10 | 1123.5340 | 562.2706 | | | 1105.5234 | 553.2654 | Y | 466.2772 | 233.6423 | 449.2507 | 225.1290 | | | 3 |
| 11 | 1279.6351 | 640.3212 | 1262.6086 | 631.8079 | 1261.6245 | 631.3159 | R | ***303.2139*** | 152.1106 | 286.1874 | 143.5973 | | | 2 |
| 12 | | | | | | | K | 147.1128 | 74.0600 | 130.0863 | 65.5468 | | | 1 |

Supplemental FIG. 1B

### MATRIX SCIENCE Mascot Search Results

#### Peptide View

MS/MS Fragmentation of **MALYVEIATYR**
Found in **gi|71011081** in **NCBInr**, mutant desmin [Homo sapiens]

Match to Query 3387: 1424.815448 from(713.415000,2+) scans(3894) rtinseconds(2076.9)
Title: function=6 process=0 scan=488
Data file G:\KL_XS_040424\KL_XS_040424_002.mzML

```
Monoisotopic mass of neutral peptide Mr(calc): 1424.6411
Fixed modifications: Propionamide (C) (apply to specified residues or termini only)
Variable modifications:
M1     : Oxidation (M), with neutral losses 63.9983(shown in table), 0.0000
Y4     : Phospho (Y)
Ions Score: 43  Expect: 1.9
Matches : 23/116 fragment ions using 51 most intense peaks   (help)
```

| # | b | $b^{++}$ | $b^0$ | $b^{0++}$ | Seq. | y | $y^{++}$ | y* | $y^{*++}$ | $y^0$ | $y^{0++}$ | # |
|---|---|---|---|---|---|---|---|---|---|---|---|---|
| 1 | 84.0444 | 42.5258 | | | M | | | | | | | 11 |
| 2 | **155.0815** | 78.0444 | | | A | ***1278.6130*** | 639.8101 | 1261.5864 | 631.2969 | 1260.6024 | 630.8048 | 10 |
| 3 | **268.1656** | 134.5864 | | | L | ***1207.5759*** | 604.2916 | 1190.5493 | 595.7783 | 1189.5653 | 595.2863 | 9 |
| 4 | 511.1952 | 256.1013 | | | Y | ***1094.4918*** | 547.7495 | 1077.4653 | 539.2363 | 1076.4812 | 538.7443 | 8 |
| 5 | 610.2636 | 305.6355 | | | V | ***851.4621*** | 426.2347 | 834.4356 | 417.7214 | 833.4516 | 417.2294 | 7 |
| 6 | 739.3062 | 370.1568 | 721.2957 | 361.1515 | E | ***752.3937*** | 376.7005 | 735.3672 | 368.1872 | 734.3832 | 367.6952 | 6 |
| 7 | **852.3903** | 426.6988 | 834.3797 | 417.6935 | I | ***623.3511*** | 312.1792 | 606.3246 | 303.6659 | 605.3406 | 303.1739 | 5 |
| 8 | 923.4274 | 462.2173 | 905.4168 | 453.2121 | A | ***510.2671*** | 255.6372 | 493.2405 | 247.1239 | 492.2565 | 246.6319 | 4 |
| 9 | 1024.4751 | 512.7412 | 1006.4645 | 503.7359 | T | ***439.2300*** | 220.1186 | 422.2034 | 211.6053 | 421.2194 | 211.1133 | 3 |
| 10 | 1187.5384 | 594.2728 | 1169.5279 | 585.2676 | Y | ***338.1823*** | 169.5948 | 321.1557 | 161.0815 | | | 2 |
| 11 | | | | | R | ***175.1190*** | 88.0631 | 158.0924 | 79.5498 | | | 1 |

### Supplemental FIG. 1C

Matched peptides shown in ***bold red***.

```
  1 MSQAYSSSQR VSSYRRTFGG APGFPLGSPL SSPVFPRAGF GSKGSSSSVT
 51 SRVYQVSRTS GGAGGLGSLR ASRLGTTRTP SSYGAGELLD FSLADAVNQE
101 FLTTRTNEKV ELQELNDRFA NYIEKVRFLE QQNAALAAEV NRLKGREPTR
151 VAELYEEELR ELRRQVEVLT NQRARVDVER DNLLDDLQRL KAKLQEEIQL
201 KEEAENNLAA FRADVDAATL ARIDLERRIE SLNEEIAFLK KVHEEEIREL
251 QAQLQEQQVQ VEMDMSKPDL TAALRDIRAQ YETIAAKNIS EAEEWYKSKV
301 SDLTQAANKN NDALRQAKQE MMEYRHQIQS YTCEIDALKG TNDSLMRQMR
351 ELEDRFASEA SGYQDNIARL EEEIRHLKDE MARHLREYQD LLNVKMALDV
401 EIATYRKLLE GEESRINLPI QTYSALNFRE TSPEQRGSEV HTKKTVMIKT
451 IETRDGEVVS EATQQQHEVL
```

Matched peptides shown in ***bold red***.

```
  1 MSQAYSSSQR VSSYRRTFGG APGFPLGSPL SSPVFPRAGF GSKGSSSSVT
 51 SRVYQVSRTS GGAGGLGSLR ASRLGTTRTP SSYGAGELLD FSLADAVNQE
101 FLTTRTNEKV ELQELNDRFA NYIEKVRFLE QQNAALAAEV NRLKGREPTR
151 VAELYEEELR ELRRQVEVLT NQRARVDVER DNLLDDLQRL KAKLQEEIQL
201 KEEAENNLAA FRADVDAATL ARIDLERRIE SLNEEIAFLK KVHEEEIREL
251 QAQLQEQQVQ VEMDMSKPDL TAALRDIRAQ YETIAAKNIS EAEEWYKSKV
301 SDLTQAANKN NDALRQAKQE MMEYRHQIQS YTCEIDALKG TNDSLMRQMR
351 ELEDRFASEA SGYQDNIARL EEEIRHLKDE MARHLREYQD LLNVKMALYV
401 EIATYRKLLE GEESRINLPI QTYSALNFRE TSPEQRGSEV HTKKTVMIKT
451 IETRDGEVVS EATQQQHEVL
```

Supplemental FIG. 2A

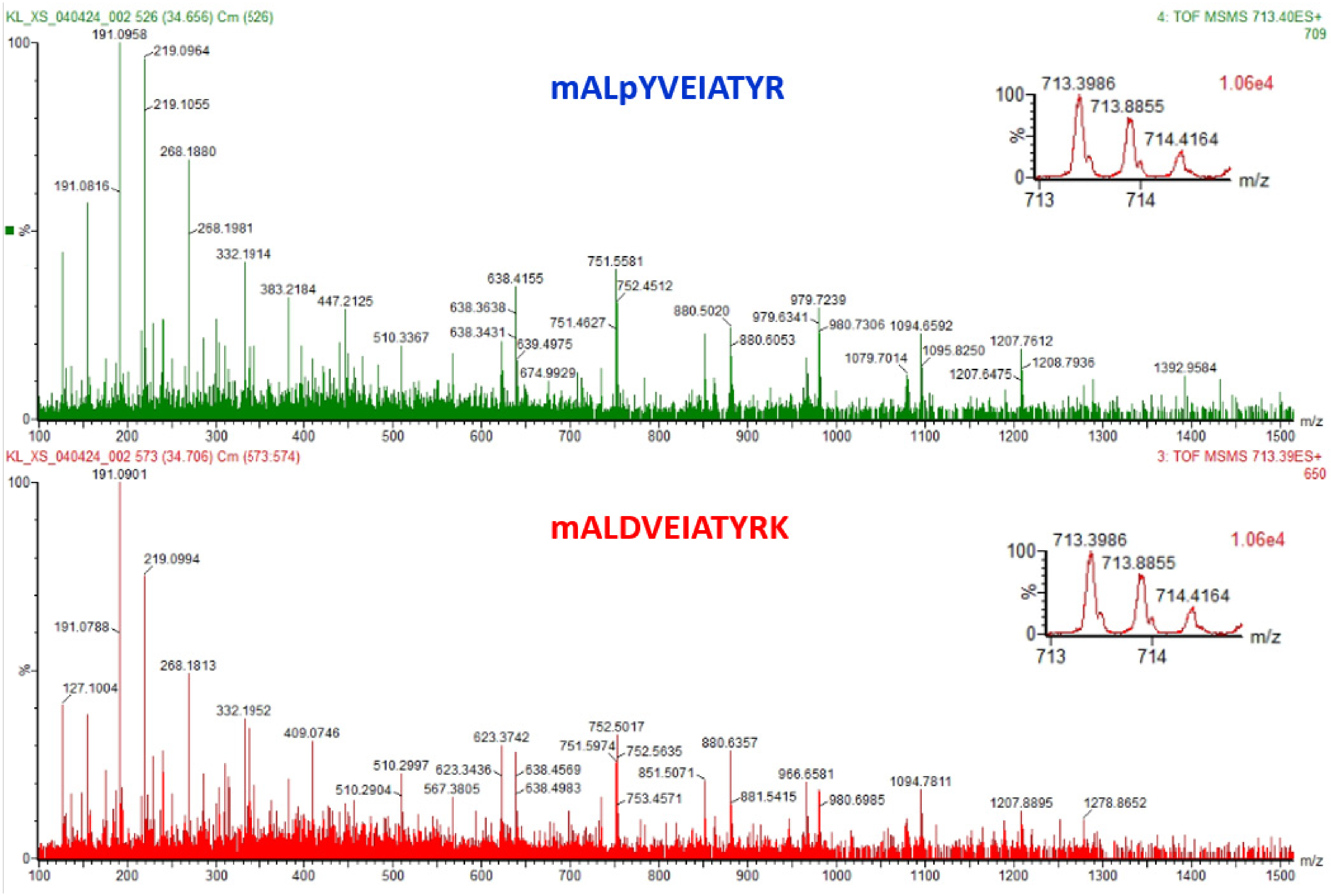

### Supplemental FIG. 2B

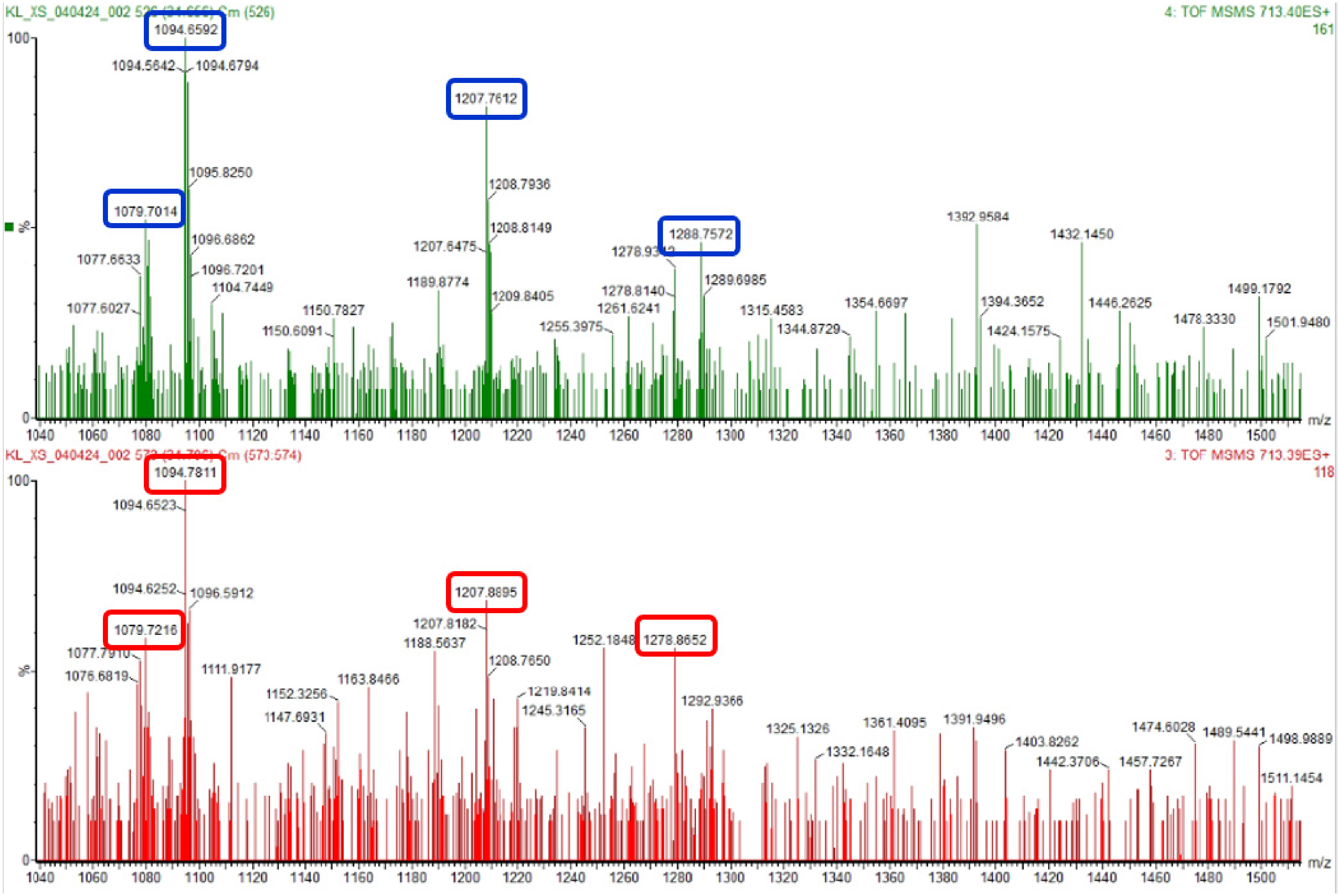

### Supplemental FIG. 2C

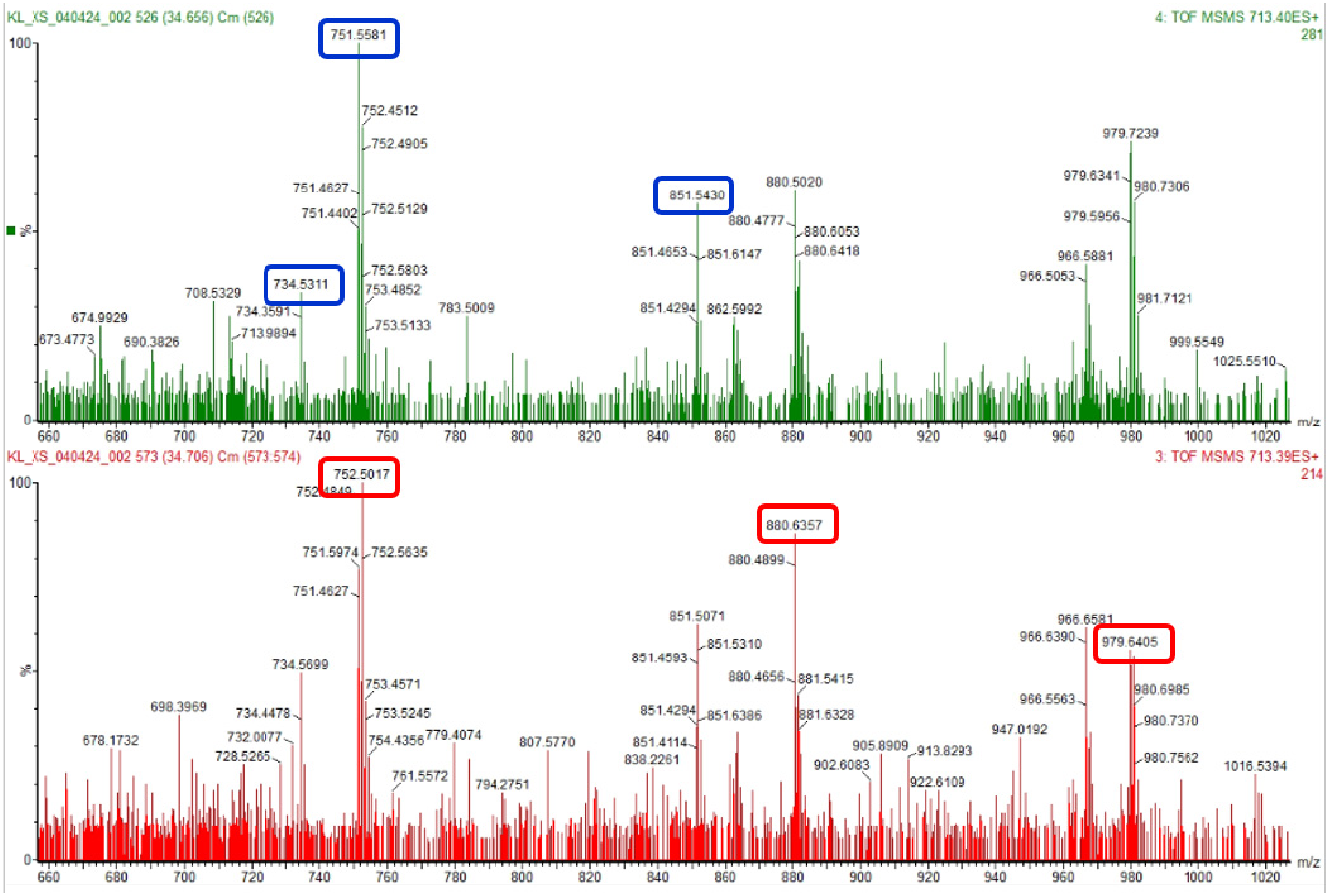

### Supplemental FIG. 2D

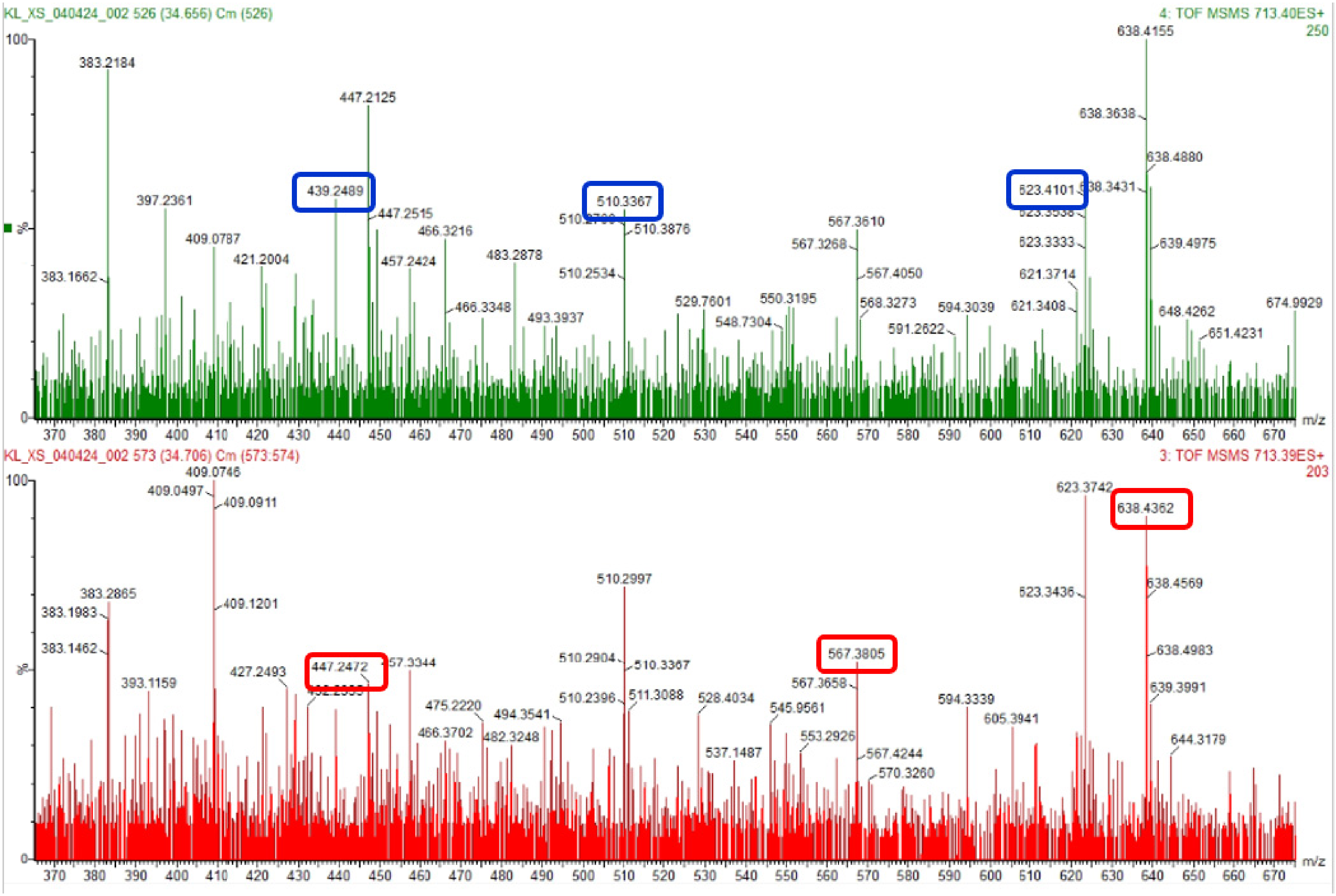

### Supplemental FIG. 2E

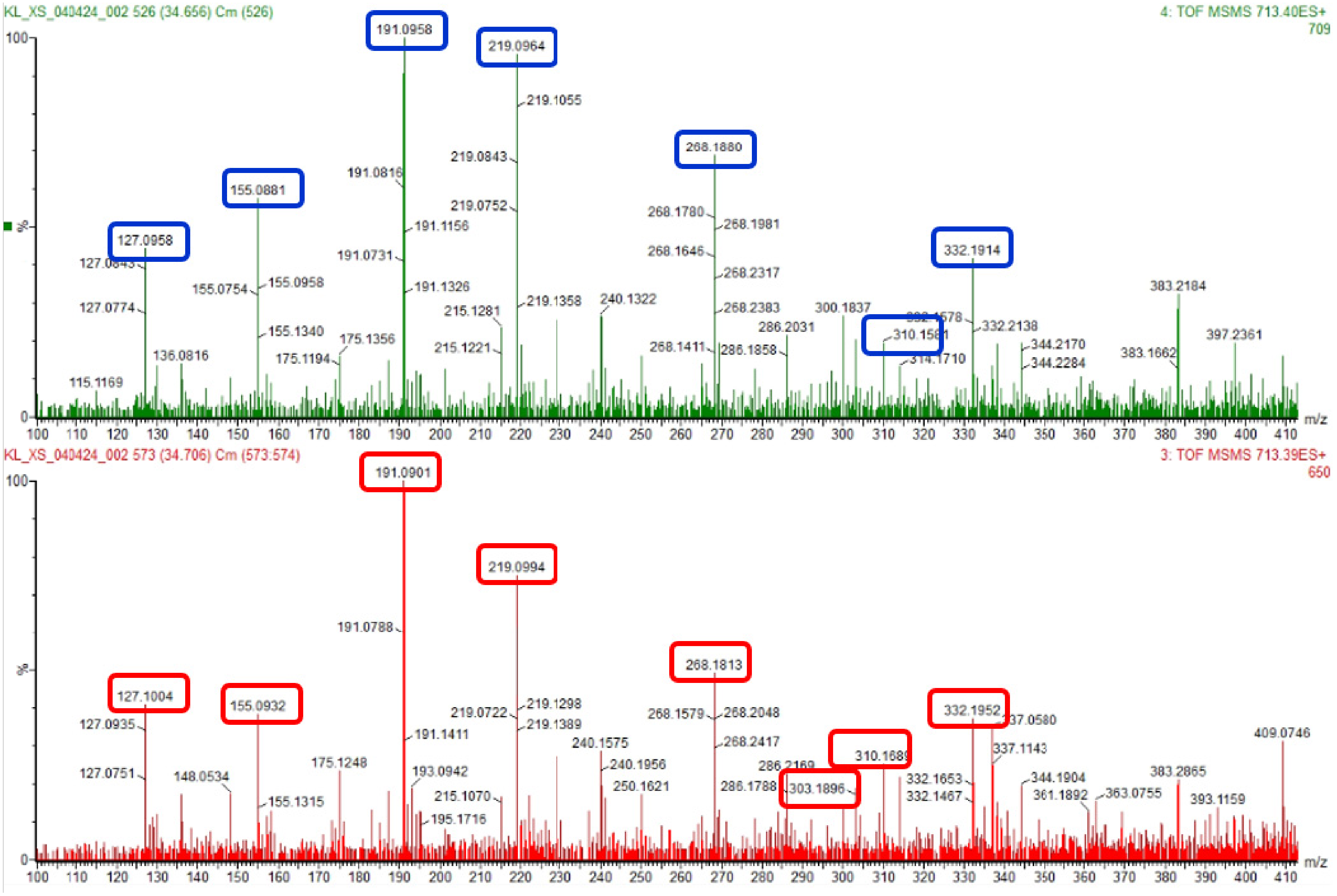

Supplemental FIG. 3A

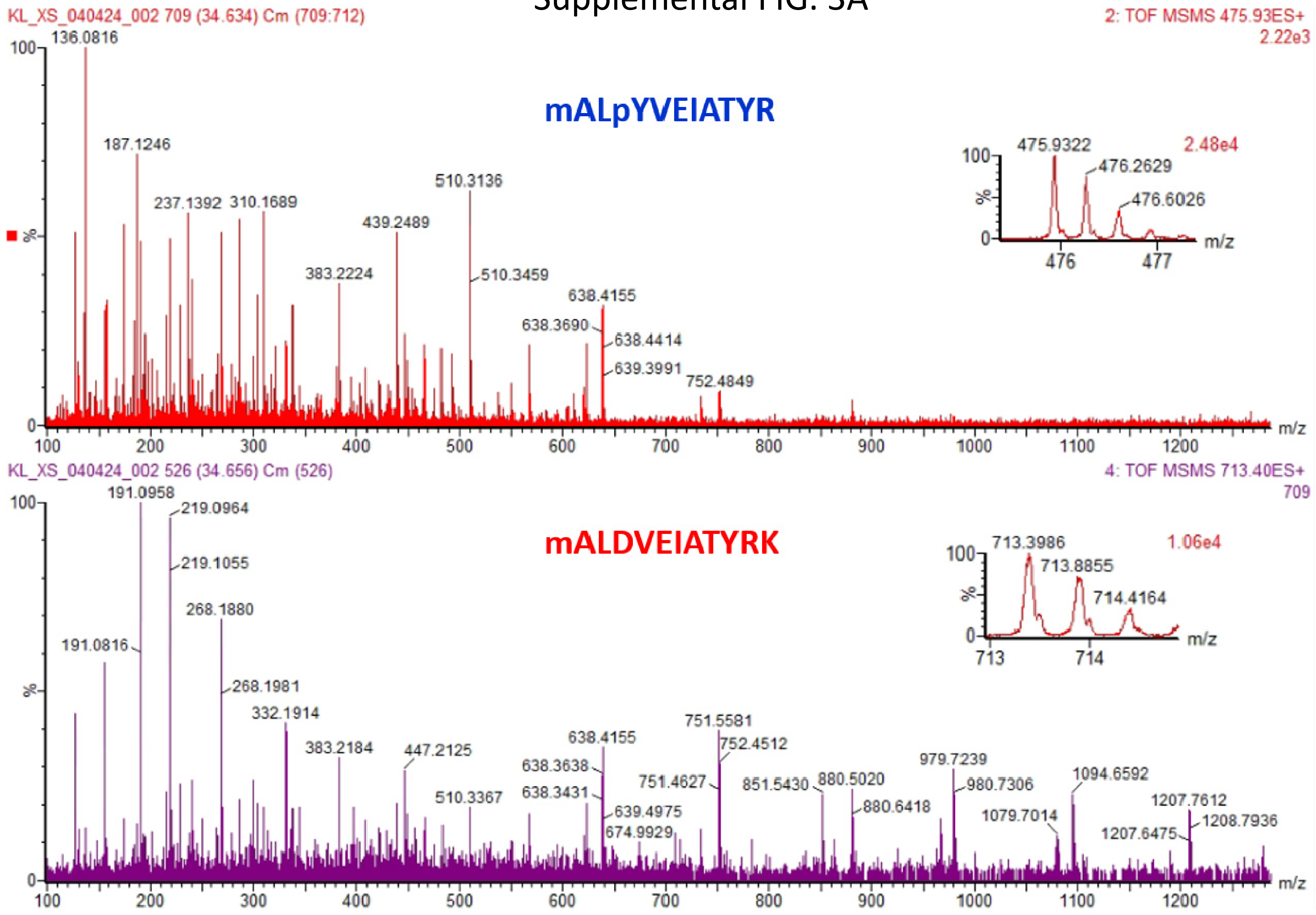

### Supplemental FIG. 3B

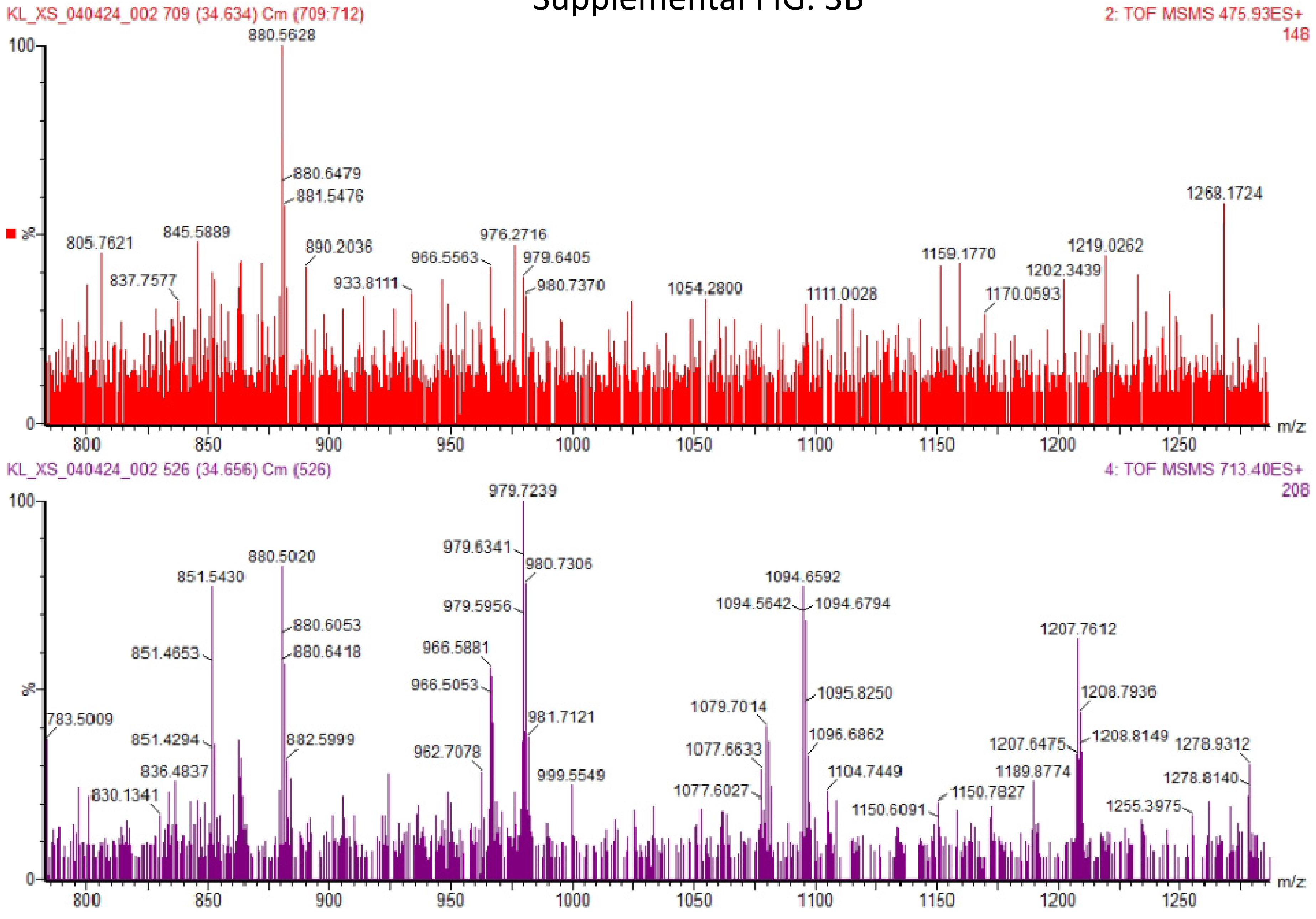

### Supplemental FIG. 3C

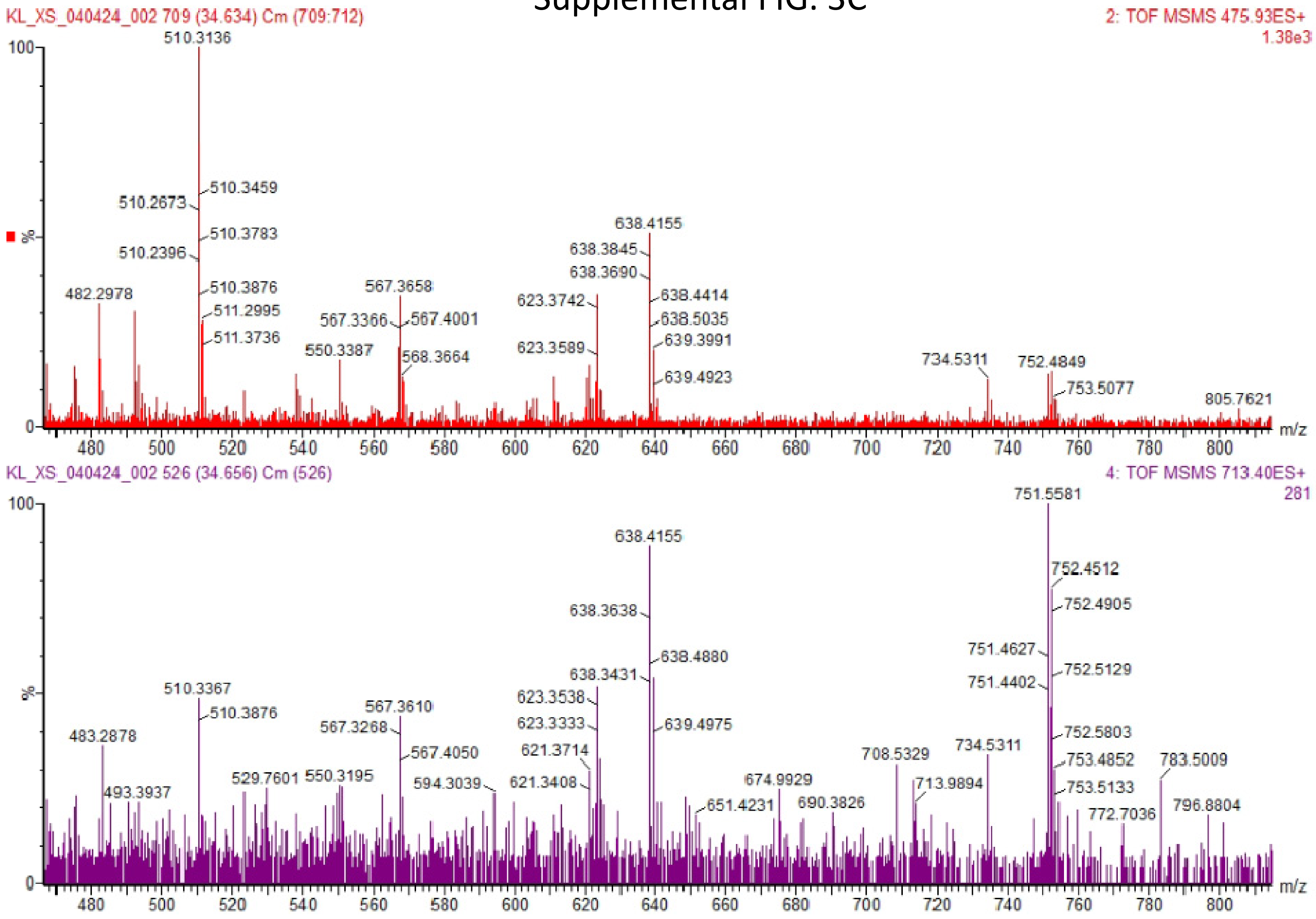

### Supplemental FIG. 3D

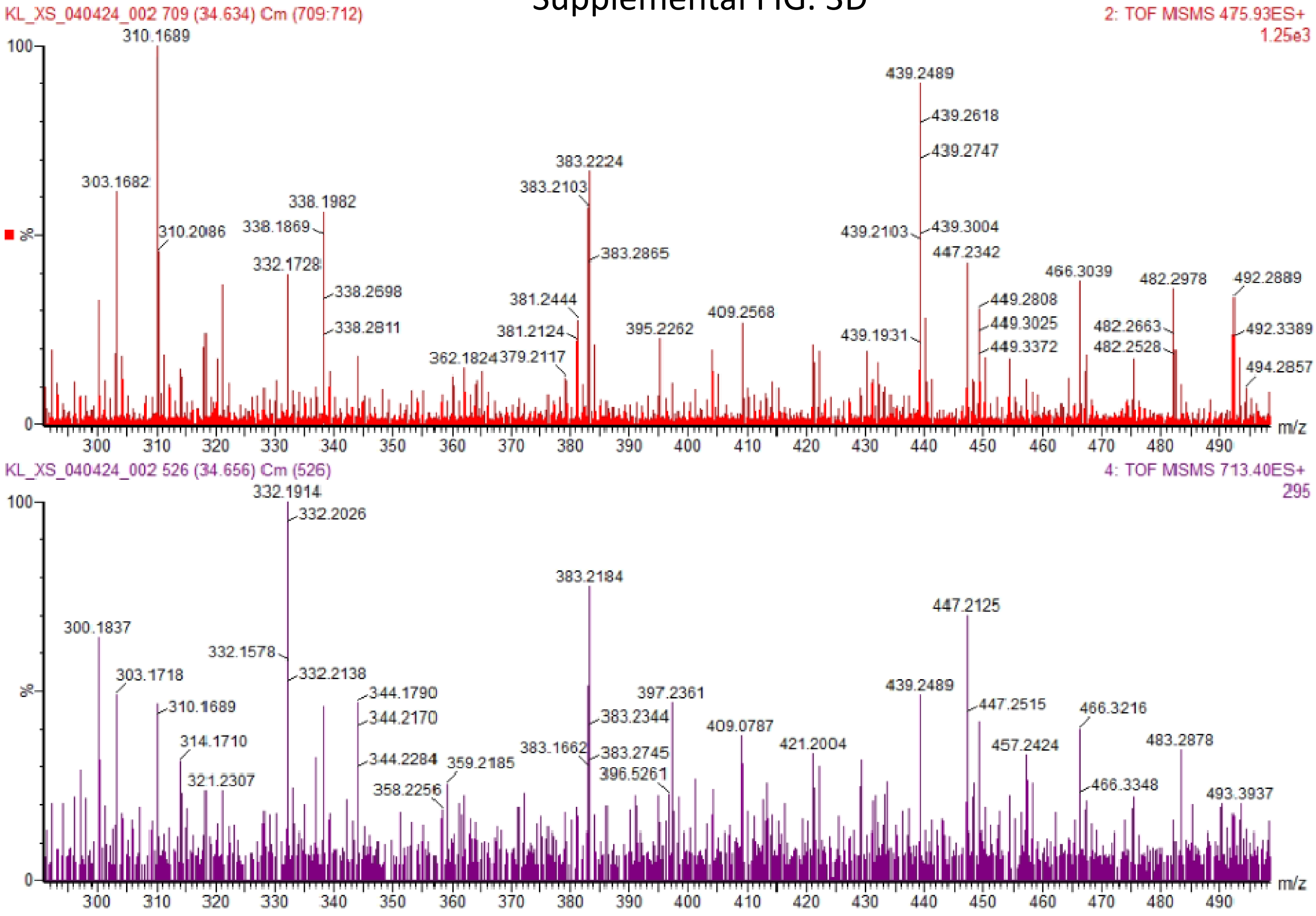

Supplemental FIG. 3E

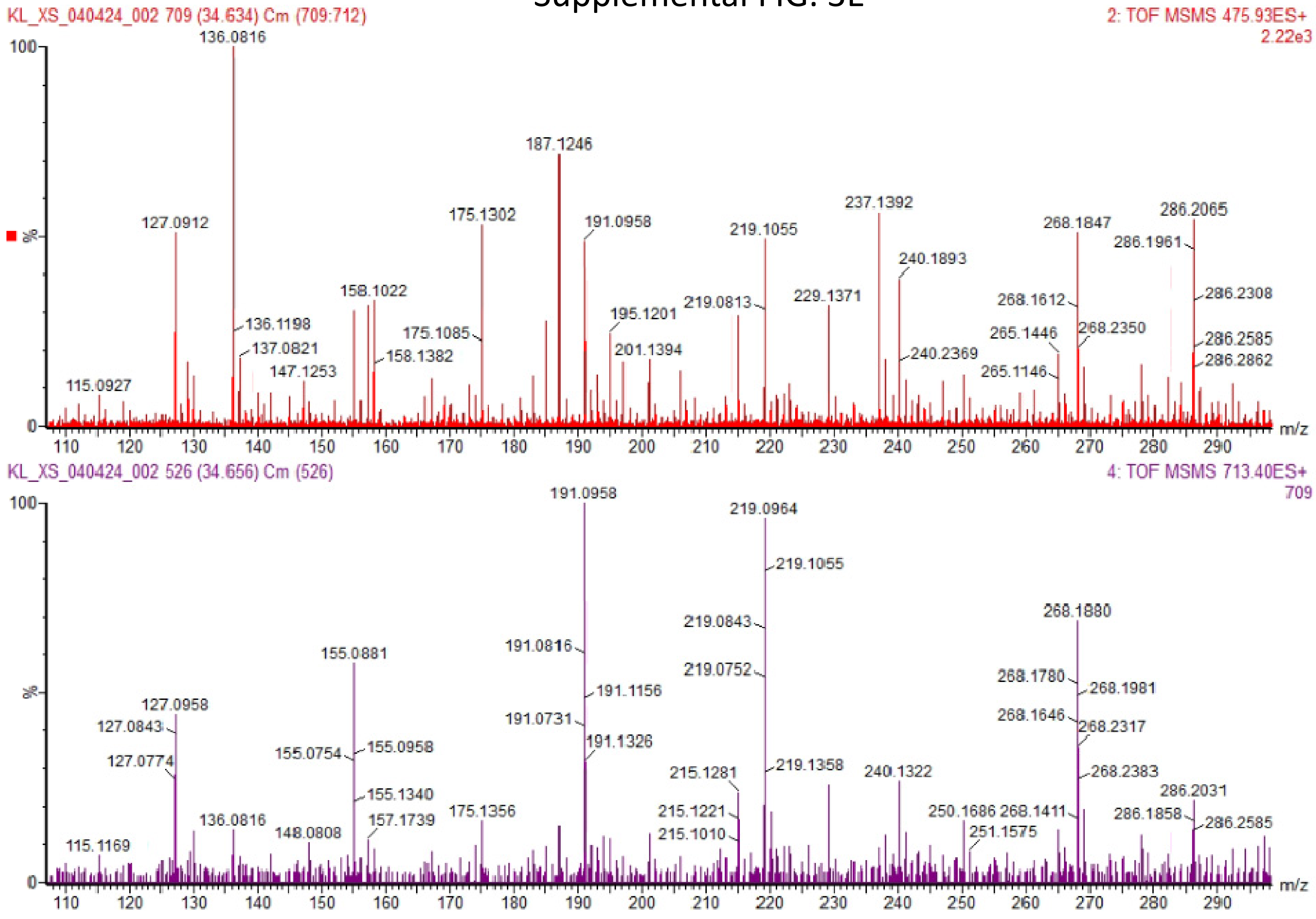
